# DFG-1 residue controls inhibitor binding mode and affinity providing a basis for rational design of kinase inhibitor selectivity

**DOI:** 10.1101/2020.07.01.182642

**Authors:** Martin Schröder, Alex N. Bullock, Oleg Federov, Franz Bracher, Apirat Chaikuad, Stefan Knapp

**Affiliations:** Institute of Pharmaceutical Chemistry, Goethe-University Frankfurt, Max von Lauestraße 9 60438 Frankfurt, Germany; Structural Genomics Consortium, Buchmann Institute for Molecular Life Sciences (BMLS), Goethe-University Frankfurt, Max von Lauestraße 15, 60438 Frankfurt, Germany; German translational cancer network (DKTK), Frankfurt/Mainz, site, 60438 Frankfurt, Germany; Structural Genomics Consortium, Nuffield Department of Medicine, University of Oxford, Oxford, OX3 7DQ, UK; Department of Pharmacy, Center for Drug Research, Ludwig-Maximilians University Munich, 81377 Munich, Germany

**Author notes:** Authors for correspondence: Stefan Knapp:, Apirat Chaikuad.

**Keywords:** kinase inhibitors, inhibitor selectivity, structure-based design, DFG-1 residue, xDFG

## Abstract

Selectivity remains a challenge for ATP-mimetic kinase inhibitors, an issue that may be overcome by targeting unique residues or binding pockets. However, to date only few strategies have been developed. Here we identify that bulky residues located N-terminal to the DFG motif (DFG-1) represent an opportunity for designing highly selective inhibitors with unexpected binding modes. We demonstrate that several diverse inhibitors exerted selective, non-canonical binding modes that exclusively target large hydrophobic DFG-1 residues present in many kinases including PIM, CK1, DAPK and CLK. Using the CLK family as a model, structural and biochemical data revealed that the DFG-1 valine controlled a non-canonical binding mode in CLK1, providing a rational for selectivity over the closely-related CLK3 which harbors a smaller DFG-1 alanine. Our data suggests that targeting the restricted back pocket in the small fraction of kinases that harbor bulky DFG-1 residues offers a versatile selectivity filter for inhibitor design.

## INTRODUCTION

Protein kinases constitute one of the largest protein families in the human genome consisting of more than 500 members^1^. Their tightly controlled catalytic activity plays a crucial role in the regulation of cellular signaling that orchestrates most cellular processes^2^. Alteration of kinase functions due to mutations or overexpression often leads to pathogenesis in a large and diverse range of diseases. Thus, targeting protein kinases offers opportunities for the development of new medicines. Many kinases are therefore currently targeted in drug development programs which have resulted in 52 FDA approved small molecule drugs^3–4^. However, most kinase inhibitors have been developed only for oncology applications. The development of highly selective inhibitors, which represents a tremendous challenge would be required for non-oncology applications.

The development of ATP competitive small molecule inhibitors for kinases has typically utilized two main binding modes. Type-I inhibitors target the active state and are anchored to the kinase hinge region by main chain hydrogen bonds that connect the two large kinase lobes. Type-II inhibitors often also interact with the kinase hinge but they target an inactive conformation characterized by reorientation of the tripeptide ‘DFG’ motif (DFG-out) that creates a large pocket accessible to ATP mimetic inhibitors^5^. Allosteric inhibitors (type-III and type-IV) have additionally been developed that are usually highly selective. However the paucity of rational design strategies has resulted in comparably few potent inhibitors so far^5–6^. Since protein kinases share in common the ATP binding pocket, achieving necessary selectivity for type-I and type-II inhibitors is a challenging task. The discovery of the DFG-out conformation in the ABL1 crystal structure in complex with the first approved kinase drug imatinib (gleevec) was met with great interest as the structure suggested that targeting this conformational change may provide a rational for the development of more selective kinase inhibitors^7^. Indeed, the type-II binding mode of imatinib is the main structural reason for its selectivity against the closely related SRC kinases. However, more comprehensive studies revealed that the DFG-out conformation can be adopted by many kinases, providing not a rational per se for selective inhibitor development^8^.

A few kinases have demonstrated other conformational changes that may lead to induced-fit pockets peripheral to the ATP binding site exploitable for the development of highly selectivity type-I inhibitors. This is exemplified by SCH772984, which binds a unique P-loop pocket in ERK1/2^9^. A rare glycine residue that allows flexibility in the hinge backbone has also been exploited for the development of skepinone-L and other p38 inhibitors^10–12^. Furthermore, targeting a unique back pocket peripheral to the αC and DFG motif offers a rationale for the high selectivity of cyclopropylethyl-substituted PI3Kγ inhibitors^13^. In addition, covalent targeting cysteine residues offers an additional strategy for the development of highly selective irreversible inhibitors^14^.

Conserved residues lining the ATP binding site can also provide a determinant controlling protein kinase sensitivity to inhibitors. One key position in the kinase ATP site is the gatekeeper residue located at the beginning of the hinge region. Small residues at this position are present in a small proportion of kinases in the human kinome and open a hydrophobic back pocket that has been exploited in the development of many kinase inhibitors, including imatinib and erlotinib drugs^8, 15–18^. The important role of the gatekeeper is highlighted by mutations at this position which confer kinase resistance to cancer drugs that target the gatekeeper back pocket^19–21^.

Here we focused on a so far little explored residue position in the kinase active site that seems to be important for inhibitor selectivity and unexpected binding modes. This critical position is located N-terminal to the DFG motif (named here DFG-1; known also as xDFG), and typically harbors a small residue. Nevertheless, larger hydrophobic amino acids, such as valine, isoleucine and leucine, are observed in a few kinases. We observed that the presence of such bulky residues in this position favored unusual binding modes and often conferred selectivity for inhibitors of this group of kinases. In addition, we were able to show that amino acid substitution at the DFG-1 position could modulate inhibitor affinities and binding modes, providing a mechanism for subfamily selectivity within the highly conserved CLKs (Cdc2 like kinases) of which all members, except CLK3 contain large hydrophobic residues in DFG-1. The presence of large DFG-1 residues in several potential drug targets such as PIM kinases, CK1 and DAPK3 offers a rational for selective inhibitor development targeting this unusual residue variation.

## RESULTS

### Unexpected similar selectivity pattern for diverse kinase inhibitors

Selectivity of kinase inhibitors is typically achieved by moderate to high degrees of chemical decorations that target unique features within the target proteins. Analysis of melting temperature shift (ΔTm) assay data from multiple kinase systems identified a set of diverse inhibitors with limited decorations that exhibited markedly high selectivity (Figure 1). These included the imidazopyridazine K00135 first reported as a PIM1 inhibitor^22^ (**1**), the benzothiazole TG003 reported as a CLK inhibitor^23–24^ (**2**), the β-carboline KH-CARB13^25^ (**3**) and the benzisoxazole-pyrimidine-2-amine also reported to target PIM kinases^26^ (**4**) (Figure 1A-B and Supplementary Table 1). The furo[3,2-b]pyridine MU1210 (**5**) presented another example with even fewer off-targets and was recently developed as a chemical probe for CLK kinases (Figure 1C)^27^.

**Figure 1.**
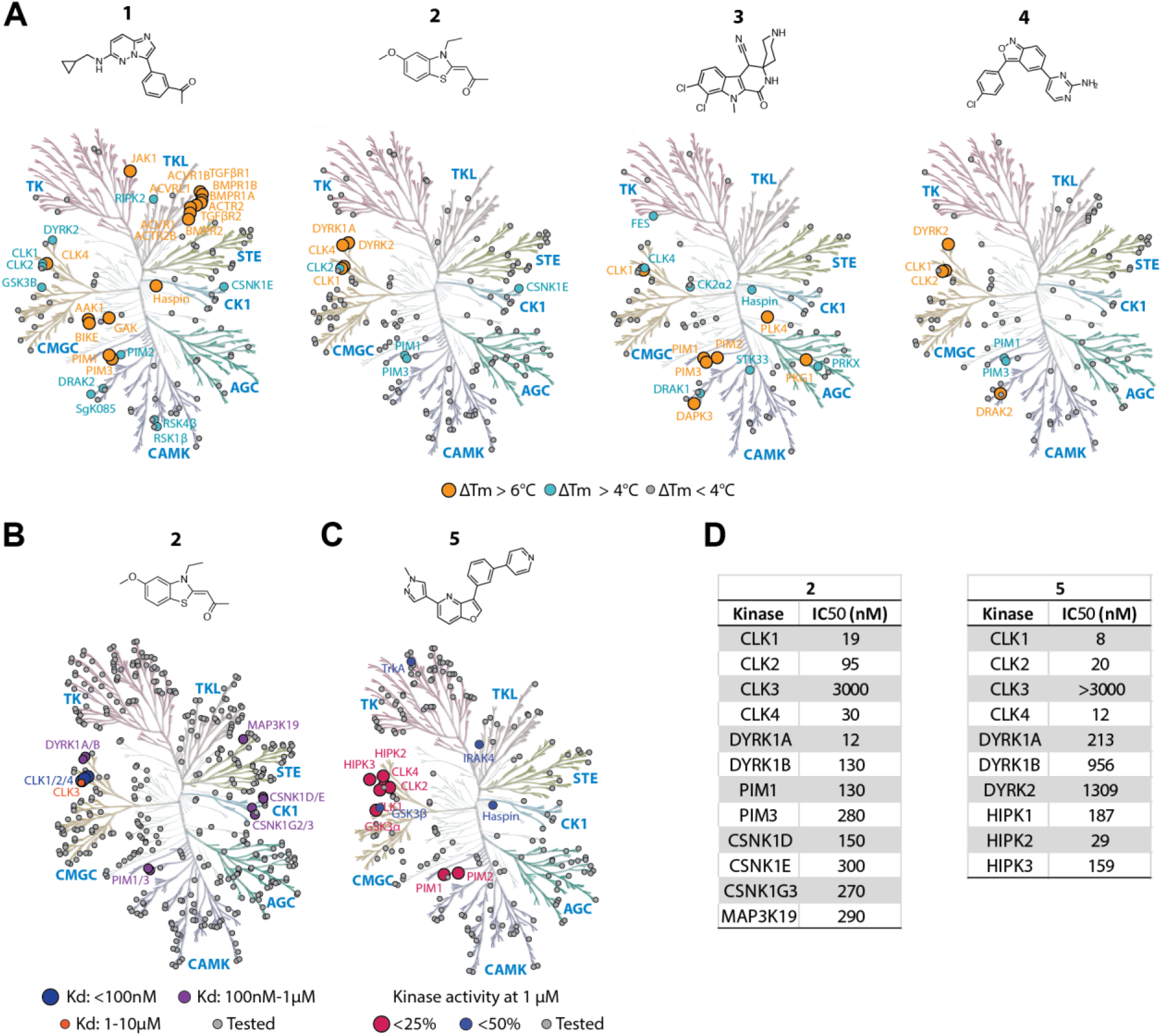
A common selectivity profile of diverse kinase inhibitors. Kinome-tree representation of the selectivity profiles of compound **1-5** measured by **(A)** our melting temperature shift (ΔTm) assays or **(B, C)** published kinome-wide profiling data^24, 27^. Gray dots indicate kinases in the assays, and colored dots demonstrates binding potencies as indicated in the schemes in each panel. **D)** IC_50_s of **2** and **5** for selected kinases^27–28^.

Of particular note was a set of diverse kinases similarly targeted by these inhibitors. This was surprising considering the structural diversity of the inhibitors. Kinases that were potently inhibited belonged to diverse groups and families, including the CMGC group CDC-like kinases (CLKs) and dual specificity tyrosine-phosphorylation-regulated kinases (DYRKs), the CAMK group PIM kinases, death-associated protein kinases (DAPKs) and DAP kinase-related apoptosis-inducing protein kinases (DRAKs), and members of the casein kinase 1 family (CK1s). Our ΔTm binding data were corroborated by the reported nanomolar potencies of inhibitors **2** and **5** in enzyme kinetic assays (Figure 1D)^27–28^. Additional activities of **1** for some TKL kinases were also observed. Thus, the selectivity data suggested a potential common mechanism responsible for the unusual binding profile of these diverse inhibitors to kinases that share only weak sequence homology in their active sites.

### The DFG-1 residue modulates and stabilizes non-canonical binding modes

The kinase targets (CLK1, DYRK1A, haspin, PIM1 and DRAK2), that were inhibited by these diverse inhibitors share limited sequence identity (Supplementary figure 1) which prompted us to investigate the structural mechanism of this reoccurring selectivity pattern.

First, we wanted to understand the unique activity of **1** for TKL family members, which represented an outlier in our selectivity comparison. The structure of **1** in complex with PIM1 published previously by our group revealed an unusual binding mode^22^. The imidazopyridazine moiety of **1** was designed as a hinge binding group, yet the PIM1 complex revealed no canonical hinge interaction and instead the imidazopyridazine formed interactions within the back pocket of the ATP site including a hydrogen bond with the VIAK motif lysine (Figure 2A). In contrast, compound **1** reverted to a canonical binding mode in ACVR1 (Figure 2B), as well as CLK1 (Figure 2C), offering a structural rationale for the observed activities of this inhibitor. The non-canonical binding mode of **1** in PIM1 might be facilitated by the presence of a proline residue in the hinge region, which offer less possibilities for canonical hinge interaction for inhibitors, but instead enable the extended, curved hinge region suitable for an accommodation of the bulky acetophenone moiety. Unusual binding modes were also detected for the 1-oxo-β-carboline **3**, an inhibitor with PIM1/3 and DAPK3 dual activity^25^ (Figure 2D). The inhibitor is expected to establish a canonical ATP mimetic binding mode through bidentate hydrogen bonds between the amide oxygen and nitrogen of the 1-oxo-β-carboline ring and the hinge as shown for inhibitors of this scaffold binding to RET and CDK2^29–30^. In contrast, in the PIM1 and DAPK3 complexes, the inhibitors flipped 180 degrees to redirect the intended hinge binding moiety towards the back pocket, positioning the halogens for halogen bonds with a hinge backbone carbonyl group. Such large differences in binding modes are rarely seen for potent kinase inhibitors.

**Figure 2.**
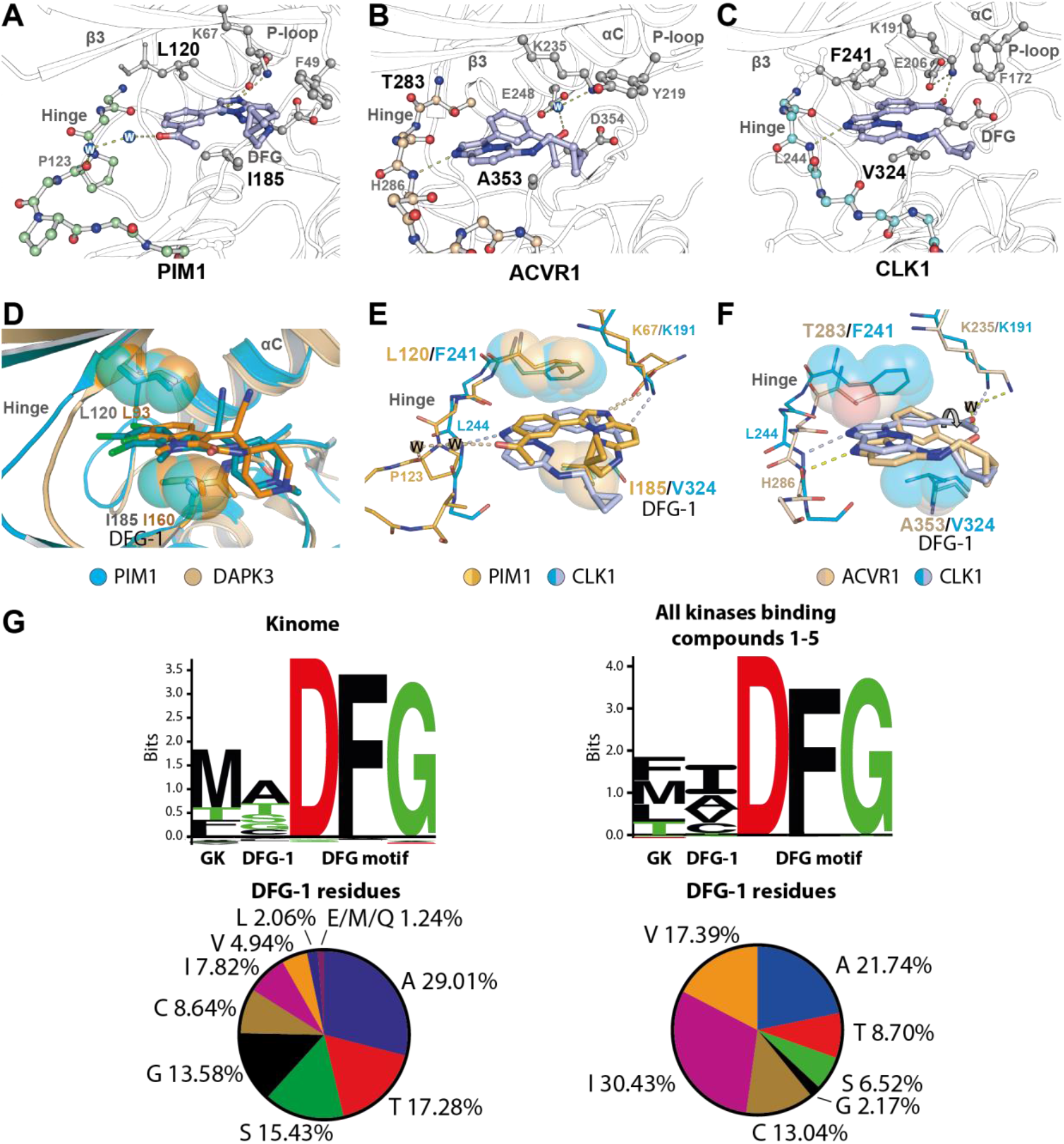
Alternative non-canonical binding modes of inhibitors stabilized by large hydrophobic DFG-1 residues. Co-crystal structures of **1** with PIM1 (**A**; pdb id 2c3i), ACVR1 (**B**; pdb id 4dym), and CLK1 (**C**; pdb id 6yta) reveals different binding modes of the inhibitor. **D**) Altered binding modes of **2** in PIM1 (pdb id 3cxw) and DAPK3 (pdb id 3bhy). Superimposition between CLK1 and PIM1 (**E**) and between CLK1 and ACVR1 (**F**) reveals potential contribution of the gatekeeper and DFG-1 residues towards the different binding modes of **1**. **G**) Distribution of amino acid compositions at the gatekeeper (GK) and DFG-1 positions across the human kinome (*left*) and among the cluster of kinases that are the targets of inhibitors **1-5** (*right*).

Further examination of the structures suggested that an additional key determinant for inhibitor binding was the combination of large hydrophobic residues at the DFG-1 and gatekeeper positions, which sandwiched aromatic ring systems in the inhibitors to stabilize the observed inverted binding modes. For instance, stabilization of the imidazopyridazine and acetophonone of **1** in the back pocket of PIM1 and CLK1, respectively, was achieved through contacts with the bulkier gatekeeper and DFG-1 residues, L120/I185 in PIM1 and F241/V324 in CLK1 (Figure 2E). The combination of large DFG-1 and gatekeeper residues also constrains the size of the back pocket binding site rendering it significantly narrower. This feature is absent in ACVR1, which has smaller residues at these positions (T283/A353) and thus larger space allowing more freedom for the inhibitor evident by an out-of-plane tilt of the inhibitor (Figure 2F). This binding mode led to the lack of contacts between **1** and ACVR1 back pocket, prompting us to postulate whether its canonical binding mode was primarily the result of the smaller DFG-1 alanine.

Our sequence analyses revealed that 84% of human kinases harbor a small hydrophilic residue in the DFG-1 position while the occurrence of large hydrophobic amino acids (leucine, valine and isoleucine) is scarce (Figure 2G and Supplementary table 2). In contrast, these large branched hydrophobic residues are common among the kinases that are the targets of inhibitors **1-5**. As the kinases that contain large DFG-1 amino acids often share low sequence identities, this unusual DFG-1 residue variation may hold promise for designing selective inhibitors that are anchored in the back pocket between the gatekeeper and large hydrophobic DFG-1 residues. To investigate and establish such potential functions of the DFG-1 residue on kinase sensitivity to inhibitors, we used human CLKs as a model system and analyzed the contribution of these residues to ligand binding using site directed mutagenesis.

### The role of the DFG-1 in the CLK subfamily for inhibitor binding

CLK kinases are key regulators of RNA splicing which is often deregulated in disease^31^. All four members of the human CLK family share highly conserved structural topology^32^. However, there are some amino acid variations within their ATP binding pockets. While most are peripheral and located in solvent expose regions, the difference at the DFG-1 position is of particular significance (Figure 3A). CLK1/2/4 harbor a DFG-1 valine residue, whereas an alanine is found at this position in CLK3. The presence of this alanine coincided with the lack of significant binding of inhibitor **1-5** for CLK3 (Figure 1), prompting us to speculate that the preferred binding of these inhibitors to CLK1/2/4 was conferred by the substitution of a DFG-1 valine. To test this hypothesis, we generated two mutants, CLK1 V324A (a ‘CLK3-like’ CLK1) and CLK3 A319V (a ‘CLK1-like’ CLK3), and assessed their binding to 18 different CLK inhibitors we had available in our laboratory using thermal shift assays. Comparison between the melting temperature shifts (ΔTm) of the mutant and wild type proteins demonstrated negative and positive ΔTm differences (ΔΔTm) of ~4 °C for the CLK1 and CLK3 pairs, respectively. This was consistent with our hypothesis, suggesting a loss in inhibitor binding for the CLK1 mutant and a gain in the CLK3 mutant (Figure 3B and Supplementary table 3). The ΔTm results were in agreement with the ~3-32-fold changes in inhibitory constants (K_i_) for **2** and **3** measured in live cells using nanoBRET^33–34^ and enzyme kinetic assays^35–36^ for CLK1 and CLK3, respectively (Figure 3C). The loss of inhibitor binding affinities against the CLK1 mutant was apparent also for other inhibitors tested, except staurosporine (**6**), with more profound changes in K_i_ of ~9-109 fold observed for KH-CB19 (**7**)^37^ and GW807982X (**8**)^38–39^ (Figure 3D and Supplementary table 4). These results suggested therefore that the smaller DFG-1 alanine is likely responsible for the loss of inhibitor binding in CLK3.

**Figure 3.**
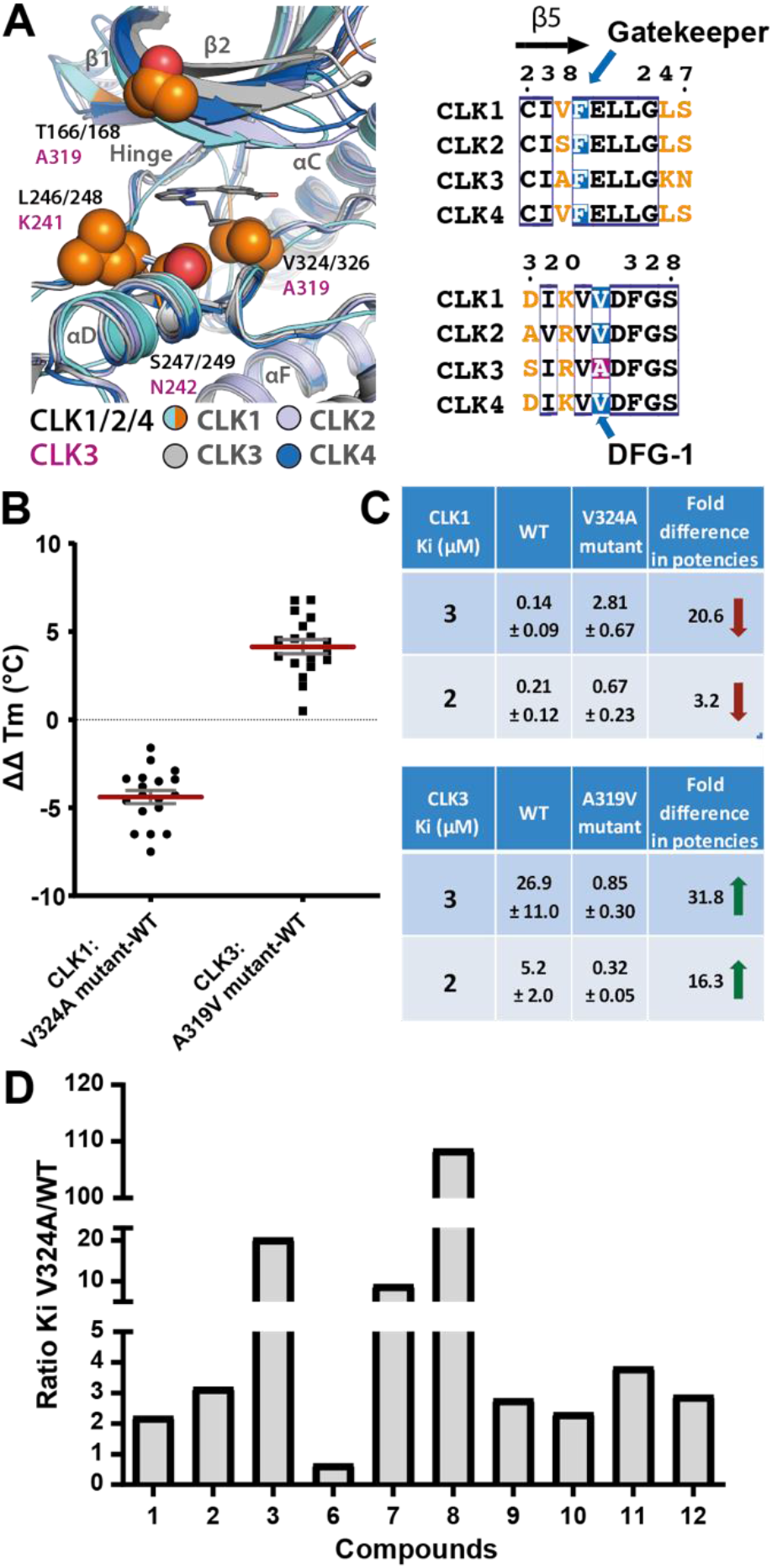
Effects of DFG-1 amino acid composition on inhibitor binding in CLK1 and CLK3. **A**) Comparative structural and sequence analyses reveal the difference within the ATP binding pockets at the DFG-1 position in the CLK subfamily, noted by a valine to alanine substitution in CLK3 (CLK1, pdb id 6yta; CLK2, pdb id 3nr9; CLK3, pdb id 6ytw; CLK4, pdb id: 6fyv^40^). **B**) Plots of melting temperature shifts (ΔTm) differences (ΔΔTm) between the mutant and wild type for 18 test compounds reveal negative and positive values for the CLK1 and CLK3 pairs, respectively (see Supplementary table 3 for ΔTm for all inhibitors). **C**) Average inhibitory constants (K_i_) for **2** and **3** measured by NanoBRET (biological duplicates) and Omnia assays (triplicates) for CLK1 and CLK3, respectively. **D**) Ratio of K_i_ between the CLK1 V324A mutant and wild type (WT) demonstrates reduction of binding potencies of most inhibitors (**9:** K00518; **10:** T3-CLK^41^; **11:** KuWal151^42^; **12:** FC162^43^).

### DFG-1 residue alters the back pocket and binding modes

To study the influence of the DFG-1 residue on inhibitor binding to CLK1 and CLK3, we selected compound **2** and **3** for more detailed structural studies to understand their distinct binding affinities for these kinases. We determined crystal structures for the inhibitor complexes of both the wild type and DFG-1 mutant kinases. Although the binding mode of **2** was not affected in all four proteins, subtle differences within the back pocket were evident. Most notable were differences in the interactions between the benzothiazole moiety and the DFG-1 residues (Figure 4A-D). A short distance of ~3.9 Å in CLK1 indicated van der Waals contacts between the inhibitor with the DFG-1 Val324, which were absent in wild type CLK3 due to its smaller DFG-1 alanine (A319). Interestingly, the DFG-1 mutations led to a reverse scenario in which the van der Waals contacts were likely diminished in the CLK1 V324A mutant, but could likely be introduced in the CLK3 A319V mutant. The loss of interaction with the DFG-1 sidechain likely explains the lower potency of **2** for CLK3 and the CLK1 V324A mutant. In addition, further examination of the structures revealed the distinct nature of the inhibitor binding cavities; a more compact pocket (~136 Å^3^) was observed in wild type CLK1 and the CLK3 mutant harboring DFG-1 valine, whereas a relatively large pocket (~217 Å^3^) was present in wild type CLK3 and the CLK1 mutant with DFG-1 alanine (Figure 4E). Isothermal calorimetry (ITC) measurements revealed minor entropic contribution, yet favorable, for the binding of **2** in both wild type CLK1 and CLK3, suggesting no significant induced change in their pocket environment. A superior enthalpy contribution observed for the **2**-CLK1 interaction was expected from the improved contacts within the restricted cavity, feasibly explaining the tighter binding of the inhibitor in this kinase (Figure 4F).

**Figure 4.**
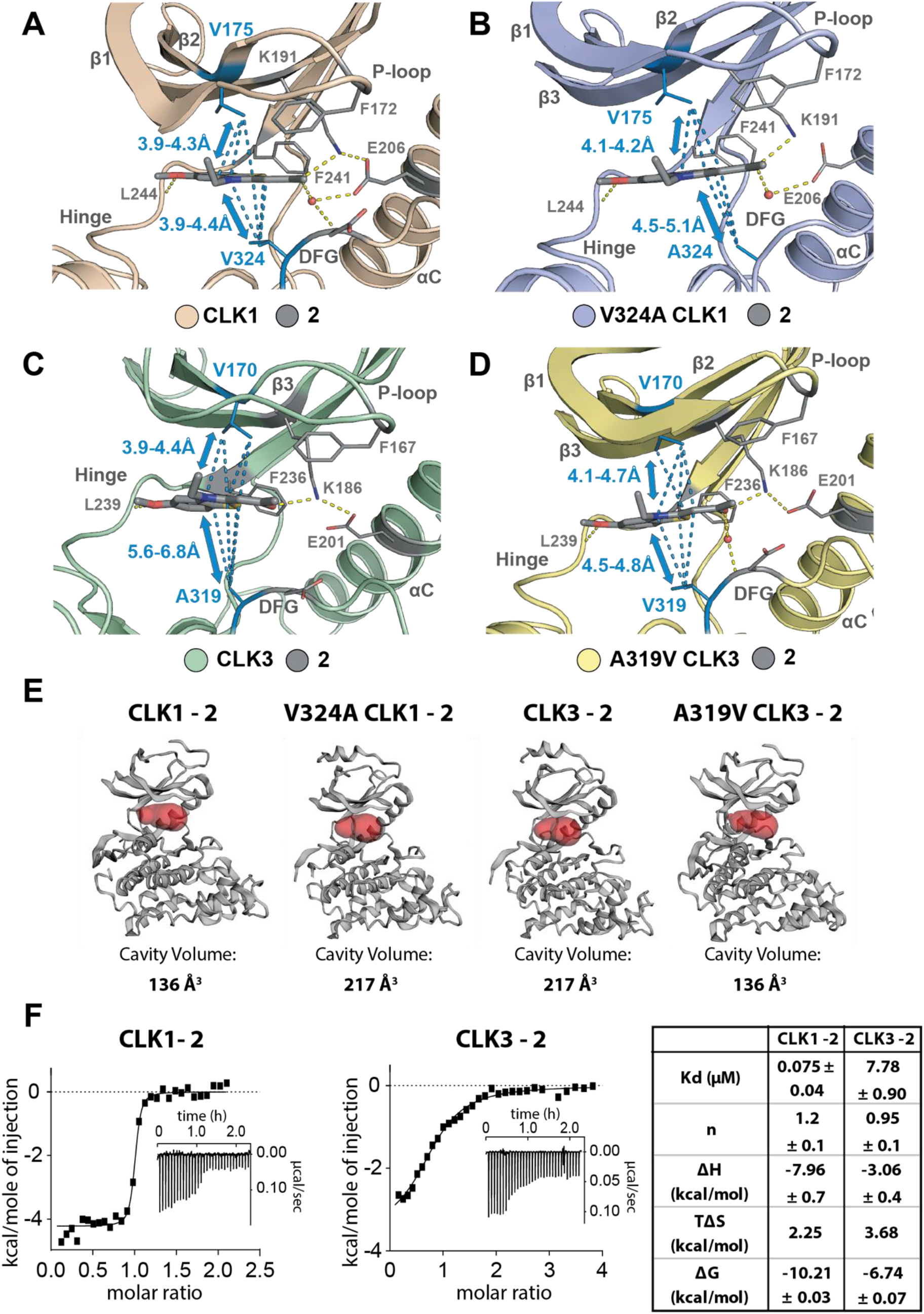
Different binding mechanisms of 2 in CLK1 and CLK3. **A-D**) The crystal structures of the complexes of **2** with wild type (CLK1: 6yte and CLK3: 6ytw) and mutant CLK1 and CLK3 (V324A CLK1: 6ytd and A319V CLK3: 6yty) demonstrate similar binding modes of the inhibitor, but different atomic distances between the β2 valine (V175 and V170 in CLK1 and CLK3, respectively) and the DFG-1 residues, suggesting different van der Waals contacts. Note the estimated coordinate errors by Luzzati plot for these structures are 0.203-0.261 Å. **E**) Pocket cavities of wild type and mutant CLK1 and CLK3 calculated for the **2-** comlexed structures. **F**) Thermodynamics of the inhibitor binding in CLK1 and CLK3 measured by ITC. Shown are integrated heat of binding, the raw isotherms (*inset*) and the average K_D_ values and thermodynamic parameters.

Surprisingly, the co-crystal structures revealed two distinct binding modes of **3** in CLK1 and CLK3 (Figure 5A-B). In the CLK1 complex, the β-carboline inhibitor formed an uncommon halogen bond with the hinge, the binding mode resembling that observed in PIM1 and DAPK3^25^. In contrast, the inhibitor flipped in CLK3 to its canonical binding mode interacting with the hinge and the solvent-exposed region. These pronounced contrast in the binding modes presumably reflected the different abilities for their back pocket to accommodate the inhibitor. The larger V324 potentially restricted the binding cavity and assisted stabilization of the β-carboline in CLK1 through van der Waals interactions, a scenario that was likely absent in the larger pocket of CLK3 with its smaller DFG-1 A319. Such a potential role for the DFG-1 residue was further supported by the CLK3 mutant structure, in which the narrowing of the back pocket by the A319V substitution led to an inverted binding mode which once again resembled that observed in wild-type CLK1 (Figure 5C). The potential DFG-1-assisted stabilization of the β-carboline in CLK1 and A319V CLK3, as well as PIM1 and DAPK3, also enabled positioning of the nitrile group in vicinity to the catalytic lysine (K191 and K186 in CLK1 and CLK3, respectively), which may lead to polar contacts explaining more favorable binding and the affinity gains in these kinases. Overall, these results established an important role for the DFG-1 residue in determining the nature of the pockets controlling kinase sensitivity. In particular, bulkier amino acids at this position may assist stabilization of high-affinity non-canonical binding mode of inhibitors, providing a basis for their selectivity.

**Figure 5.**
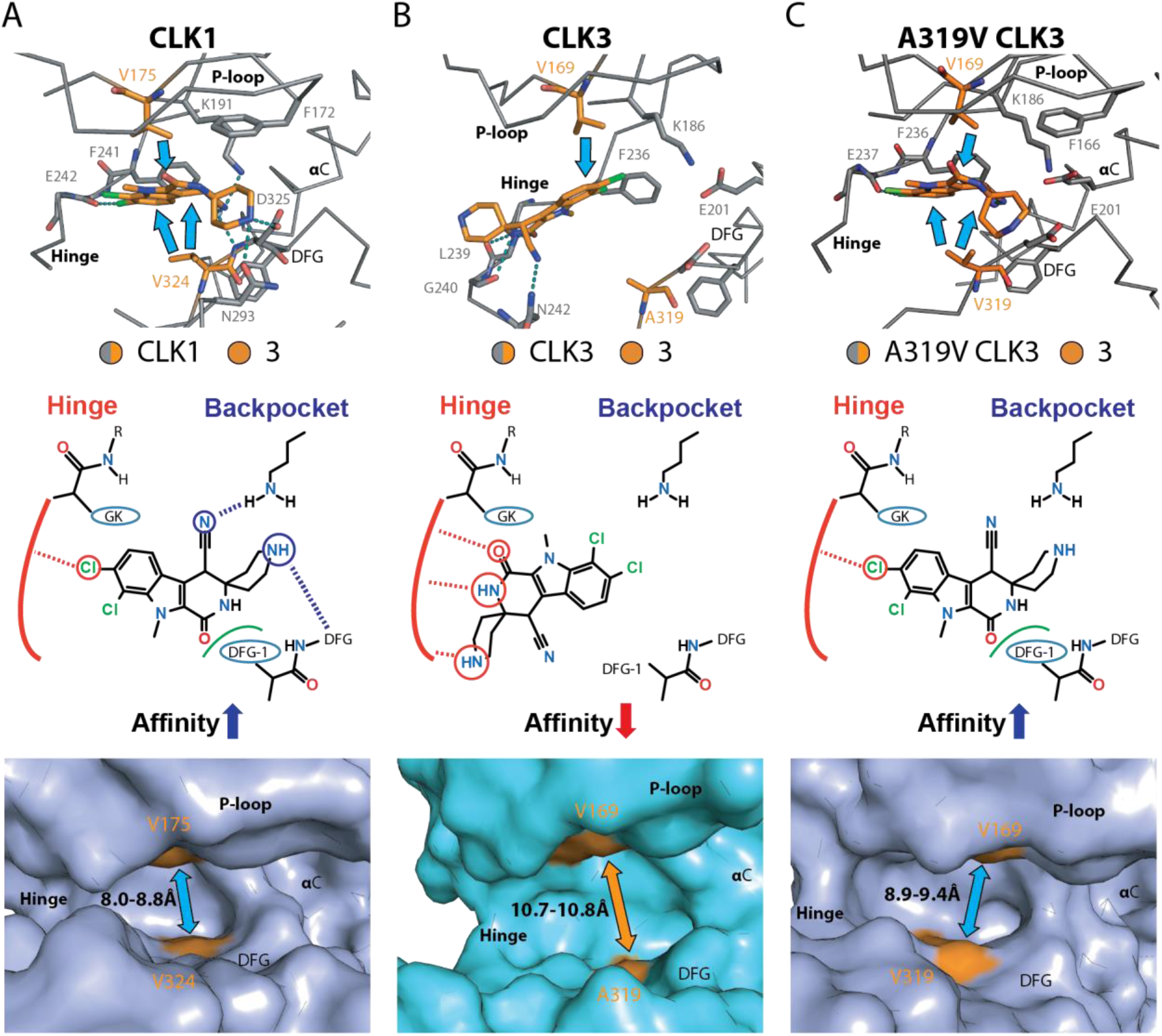
Non-canonical and canonical binding modes of 3 within the back pocket of CLK1 and CLK3. Comparison of the binding modes of **3** within CLK1 wild type (**A**; 6ytg), CLK3 wild type (**B**; 6yu1) and A319V CLK3 mutant (**C**; 6z2v). Detailed interactions of the inhibitor within the kinases are shown in the top and middle panels, while the lower panels illustrate the different pocket cavities among these kinases. Blue arrows indicate potential van der Waals contacts, while dashed lines indicate hydrogen bonds.

### Targeting the DFG-1 residue determines selectivity of CLK1/2/4 over CLK3

Our characterization showed that the restricted back pocket and the larger DFG-1 residue provided a mechanism for the high potencies of **2** and **3** for CLK1 and selectivity over CLK3. To investigate whether other inhibitors developed for this kinase family might share similar binding behavior, we profiled seven inhibitors reported previously to bind CLK1 using ΔTm assays (Figure 6A). Interestingly, all screened compounds exhibited similar behavior to **2** and **3** with binding preference for CLK1/2/4 over CLK3. Analyses of the crystal structure of CLK1 with GW807982X (**8**) and ETH1610 (**13**)^44^ revealed that these inhibitors exhibited similar binding mechanisms by tightly interacting with DFG-1 V324 (Figure 6B). Again, the CLK1 V324A mutant showed reduced inhibitor binding potency in contrast to an increase in affinities for the CLK3 A319V mutant (Figure 6B). Overall, these results demonstrate that many diverse scaffolds can exhibit such atypical binding mechanisms by targeting the bulkier DFG-1 valine residue and achieve high potency for CLK1/2/4 with selectivity over CLK3.

**Figure 6.**
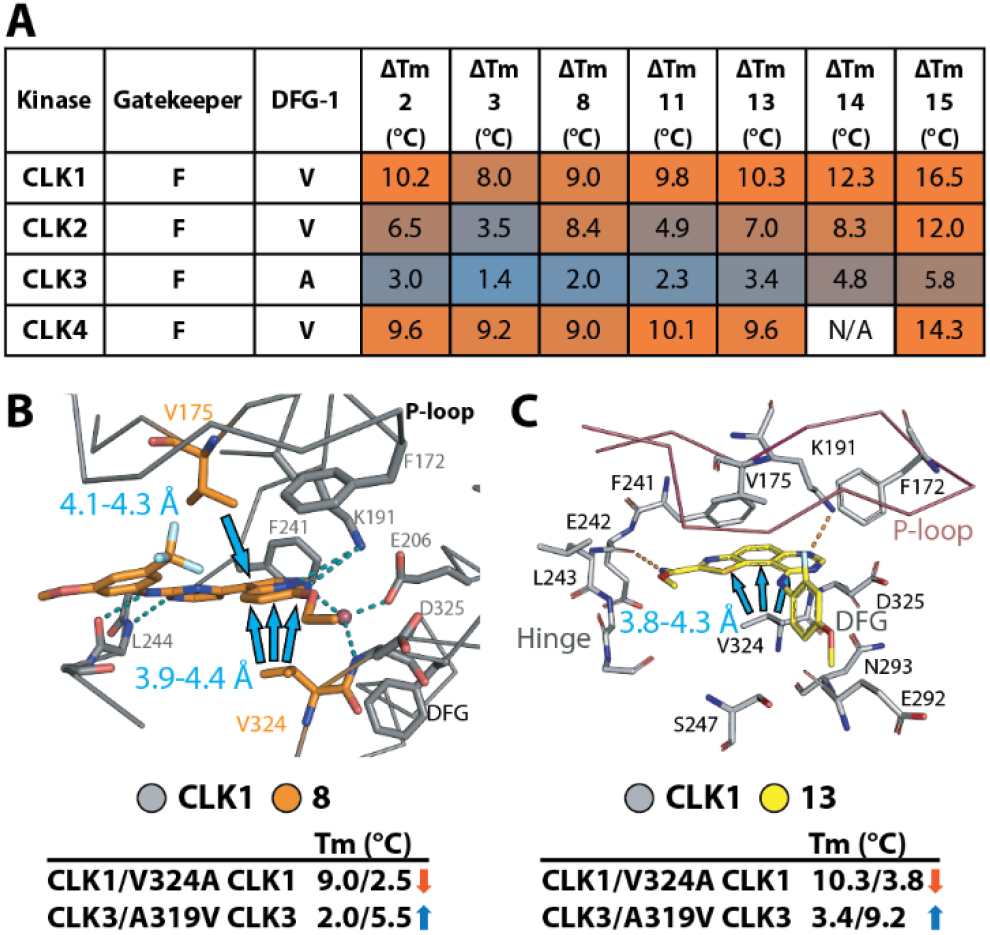
Unusual binding mechanism of targeting the DFG-1 residue for subfamily selectivity is shared by several CLK inhibitors. **A)** ΔTm for reported CLK inhibitors demonstrates selectivity over CLK3 (EHT1610 (**13**)^44^, VN412 (**14**)^27^, GW779439X (**15**)^38^). The crystal structures of CLK1 with **8** with **8** (**B**; 6zln) and **13** (**C**; 6yti) reveals the binding modes of these inhibitors likely depend on van der Waals interactions (blue arrows) with DFG-1 V324 in the back pocket. Lower panels show ΔTm for CLK1 and CLK3 wild type and mutants, indicating the changes of inhibitor binding affinities that corresponds to the different DFG-1 amino acids.

## DISCUSSION AND CONCLUSION

The rational design of kinase inhibitor with high selectivity is a challenging task. Although unique pockets peripheral to the ATP binding site may offer an opportunity to overcome this issue^9–10, 13^, the discovery of structural features present only in few kinases is rarely reported. Targeting small gatekeeper residues that are present in a small proportion of kinases in the human kinome is now widely used and has led to selective inhibitors in particular for receptor tyrosine kinases^15–17^ as well as members of the TGFβ receptor family^45^. Here we reported another alternative strategy for selective inhibitor design that makes use of bulky hydrophobic residues at the DFG-1 position present in only 15% of all human kinases. These kinases are present in most families of the human kinome and the low sequence conservation among these unrelated kinases with bulky DFG-1 can be exploited to achieve exclusive selectivity for a kinase target. We showed in this study that in the case of the CLK family the CLK3 isoform is often spared by inhibitors that make contact with the DFG-1 residue. The strategy is particularly attractive as inhibitors binding to the back pocket of kinases that harbor large DFG-1 residues often have unusual binding modes, such as compound **3**, or rely less on classical hinge interactions, such as compound **2**. The unexpected binding behavior of CLK family kinases has been reported previously^40, 46^, but the structural rationale has remained unknown. Here we demonstrated the role of the DFG-1 residue by site directed mutagenesis as well as structural insights, revealing that large residues at this DFG-1 position control the high-affinity, non-canonical binding modes that target the back pocket. This likely provides a mechanism for the enhanced selectivity of most CLK1/2/4 inhibitors over closely-related CLK3. It is interesting to note that the most selective inhibitors including the casein kinase 2 alpha 1 (CK2α1) inhibitor GO289^47^, the dual PIM/DAPK3 inhibitor HS56^48^, the selective DAPK3 inhibitor HS94 and HS184^48^ and the CLK inhibitor MU1210^27^ do not significantly interact with the hinge back bone, but rather engage contacts at the back pocket particularly with the DFG-1 residue. Such binding modes have also been noted by the KLIFS database^49^. Strong hinge binding motifs may also result in canonical hinge binding modes as observed for compound **1** binding to ACVR1 and CLK1 resulting in less favorable selectivity profiles. Thus, large and bulky DFG-1 residues offer a general anchor point for inhibitor design independent of the commonly used ATP mimetic hinge region. The improved activities and selectivity profiles of the selected inhibitors demonstrated here suggest therefore that targeting large DFG-1 residues and unusual back pockets presents an attractive general strategy for the design of new chemical libraries for many kinases.

## EXPERIMENTAL SECTION

### Compounds

All compounds used in this study were obtained commercially or from sources listed in Supplementary table 3. Purity of all compounds is > 95% as confirmed by the sources.

### Protein expression, purification and structure determination

Recombinant wild type CLK1 (H148-I484) and CLK3 (R134-T484) and their mutants, which were generated by PCR-based site directed mutagenesis, were expressed in *E. coli* and purified using the procedure described previously^32^. Crystallization of the apo kinases and their inhibitor complexes was performed using sitting vapor diffusion method at 4 °C and the conditions listed in Supplementary Table 5. Soaking was performed with 5 mM inhibitors.

For ACVR1, the kinase domain of ACVR1 R206H (residues 201-499) was subcloned into the vector pFB-LIC-Bse for baculoviral expression in Sf9 cells at a density of 2×10^6^/ml grown at 27°C, shaking at 110rpm. At 48 hours post infection cells were harvested and resuspended in 50 mM HEPES, pH 7.5, 500 mM NaCl, 5 mM imidazole, 5% glycerol, 0.1 mM TCEP supplemented with protease inhibitor set V (Calbiochem). Cells were lysed using a C5 high pressure homogenizer (Emulsiflex) then centrifuged at 21,000 rpm and 4°C for 1 hr and the supernatant recovered for purification. The protein were purified initially by Ni-affinity chromatography before being treated with TEV protease to remove the His tag. Further purification was achieved by size exclusion chromatography using a S200 HiLoad 16/60 Superdex column. The cleaved ACVR1 protein was finally passed through another Ni-affinity chromatography column before being buffered for storage in 50 mM HEPES pH 7.5, 300 mM NaCl, 10 mM DTT and 10mM L-arginine, 10 mM L-glutamate at a concentration of 10 mg/mL. Crystals of ACVR1 R206H with K00135 were grown at 20°C in a sitting drop by mixing 75 nL protein solution with 75 nL of a reservoir solution containing 1.6 M MgSO_4_ and 0.1 M MES pH 6.5.

All crystals were cryo-protected using 25% ethylene glycol, and diffraction data were collected at BESSY, Diamond Light Source and SLS. Data were processed using iMosflm^50^ or XDS^51^, and subsequently scaled using aimless^52^. Molecular replacement was performed using Phaser^53^ and the coordinates of CLK1 (pdb id 1z57)^32^, CLK3 (pdb id 2eu9)^32^ and ACVR1 (pdb id 3h9r)^54^. Model rebuilding alternated with structure refinement were performed in COOT^55^ and REFMAC^56^, respectively. Geometric correctness was validated using Molprobity^57^. Solvent accessible area was calculated using CASTp^58^ with a 2-Å cut-off radius. Data collection and refinement statistics are summarized in Supplementary Table 5, and the omitted electron density maps for the bound ligands are shown in Supplementary figure 2.

### Sequence alignment analyses

Alignment of the kinase domain sequences of 536 kinases was performed using Clustal Omega^59^. Relative occurrence of amino acids was analyzed using the P-weighted Kullback-Leibler method from which the plots were generated using SeqtoLogo v2.0^60^.

### Melting temperature shift assays

Recombinant protein at 2 μM in 10 mM HEPES, pH 7.5 and 500 mM NaCl was mixed with 10 μM inhibitors. The reaction was incubated at room temperature for 10 minutes. SyPRO orange (Invitrogen) was added at 1:1000 dilution, and the fluorescence signals corresponding to temperature-dependent protein unfolding were measured using a Real-Time PCR Mx3005p machine (Stratagene). Melting temperature shifts were calculated using the previously described method^61^.

### Omnia kinase assay

Recombinant CLK3 wild type and A319V mutant at 60 nM in reaction buffer containing 20 mM Tris, pH 7.5, 25 mM MgCl2, 10% glycerol were mixed with 40 μM Sox-modified substrate peptide (Ac-ERMRPRKRQGSVdP(Sox)G-NH2). Compounds in a serial dilution were added to the reaction mixture, and after incubation for 5 minutes at 25 °C substrate phosphorylation was initiated by addition of 0.1 mM ATP. The kinase activity was monitored by an increase in fluorescence signal using a Tecan Spark or BMG Clariostar instrument with 360 nm excitation and 460 nm emission wavelengths. Concentration-dependent changes in initial velocities were fitted using a dose-response equation in Prism, and the measured IC_50_ values were used for the inhibitory constant (K_i_) calculation using Cheng-Prussof equation and the ATP K_m_ values of 58.9 ± 20.3 and 8.6 ± 4.2 μM calculated from triplicate for the wild type and mutant, respectively.

### NanoBRET assays

Full-length CLK1 wild type and V324A mutant were subcloned into pFC32K (promega) with N- or C-terminal NanoLuc fusion, respectively. The plasmids were transfected into HEK 293T cells, which were cultured for 20 hours at 37 °C and 5% CO_2_ to allow for protein expression. The NanoBRET assays were performed as described previously^62^. In brief, after incubation cells were harvested and resuspended in Opti-MEM media, and subsequently mixed and incubated with inhibitors and 1 μM Tracer K5 (promega) for 2 hours. The mixture of NanoBRET™ Nano-Glo^®^ substrate and Extracellular NanoLuc Inhibitor (Promega) was added immediately prior to BRET luminescence measurement (450 nm for donor emission and 610 nm for acceptor emission) using a PHERAstar FSX plate reader (BMG Labtech). Milli-BRET units (mBU) calculated as a ratio of BRET signal to the overall luciferase signal were fitted using a dose-response equation in Prism. The obtained IC_50_ values were used for K_i_ calculation using a Tracer K5 K_d_ of 0.254 and 0.222 μM for the wild type and V324A mutant, respectively.

### Isothermal titration calorimetry

ITC measurements were performed in a NanoITC (TA Instruments) at 15 °C in a buffer containing 50 mM HEPES, pH 7.5, 500 mM NaCl, 50 mM arginine, 50 mM glutamate, 0.5 mM TCEP and 5% glycerol. The kinase at 130 μM was injected into the reaction cell, which contained inhibitor **2** at 15 μM. The integrated heat of titration was calculated and fitted to an independent binding model according to the manufacture’s protocol. The thermodynamic parameters, dissociation constants (KD), and stoichiometry (n) were calculated.

## Supporting Information

The Supporting information includes Figures and Tables.

## Corresponding Author

Stefan Knapp

Apirat Chaikuad

## Notes

The coordinates and structure factors of all complexes have been deposited to the protein data bank under accession codes 6yta, 6yte, 6ytd, 6ytw, 6yty, 6yu1, 6z2v, 6zln, 6yti, 4dym, 3nr9.

## ACKNOWLEDGMENT

The authors are grateful for support by the SGC, a registered charity (number 1097737) that receives funds from AbbVie, Bayer Pharma AG, Boehringer Ingelheim, Canada Foundation for Innovation, Eshelman Institute for Innovation, Genome Canada, Innovative Medicines Initiative (EU/EFPIA) [ULTRA-DD grant no. 115766], Janssen, Merck KGaA Darmstadt Germany, MSD, Novartis Pharma AG, Ontario Ministry of Economic Development and Innovation, Pfizer, São Paulo Research Foundation-FAPESP, Takeda, and Wellcome [106169/ZZ14/Z]. We are grateful to the DFG funded Collaborative Sonderforschungsbereich 1177 Autophagy (SFB1177, to MS, AC and SK) at Frankfurt University, as well as the German Translational Cancer Consortium. The authors thank staff at BESSY II, SLS and the Diamond Light Source beamline IO2 (proposal mx442) for their support during crystallographic X-ray diffraction testing and data collection. The data collection at SLS was supported by funding from the European Union’s Horizon 2020 research and innovation program under grant agreement number 730872, project CALIPSOplus.

## ABBREVIATIONS

CLK: Cdc2 like kinase
PIM: Proviral insertion in murine kinase
ACVR1: Activin receptor type I
DAPK: death-associated protein kinase
DYRK: dual specificity tyrosine-phosphorylation-regulated kinases
DRAK: DAP kinase-related apoptosis-inducing protein kinases
CK: casein kinase
RET: Proto-oncogene tyrosine-protein kinase receptor Ret
CDK2: Cyclin-dependent kinase 2
DFG motif: tripeptide Asp-Phe-Gly motif
VIAK motif: valine-isoleucine-alanine-lysine motif.

