## Supplementary information for "DFG-1 residue controls inhibitor binding mode and affinity providing a basis for rational design of kinase inhibitor selectivity"

Page

**Supplementary figure 1.** Comparative sequence analyses of the most common kinase targets of 1-5. S2

**Supplementary figure 2.** Polder difference maps at 3σ for the bound compounds S3

**Supplementary table 1.** Thermal shift results of compounds 1-4 against various kinases S4

**Supplementary table 2.** The gatekeeper and DFG-1 amino acid compositions of the kinases that interact with inhibitor **1-5.** S7

**Supplementary table 3.** ΔTm data for wild type and DFG-1-mutated CLK1 and CLK3. S8

**Supplementary table 4.** Inhibition constant (Ki) from nanoBRET assays for CLK1 wild type and V324A mutant. S9

**Supplementary table 5.** Data collection and refinement statistics. S10

**A**

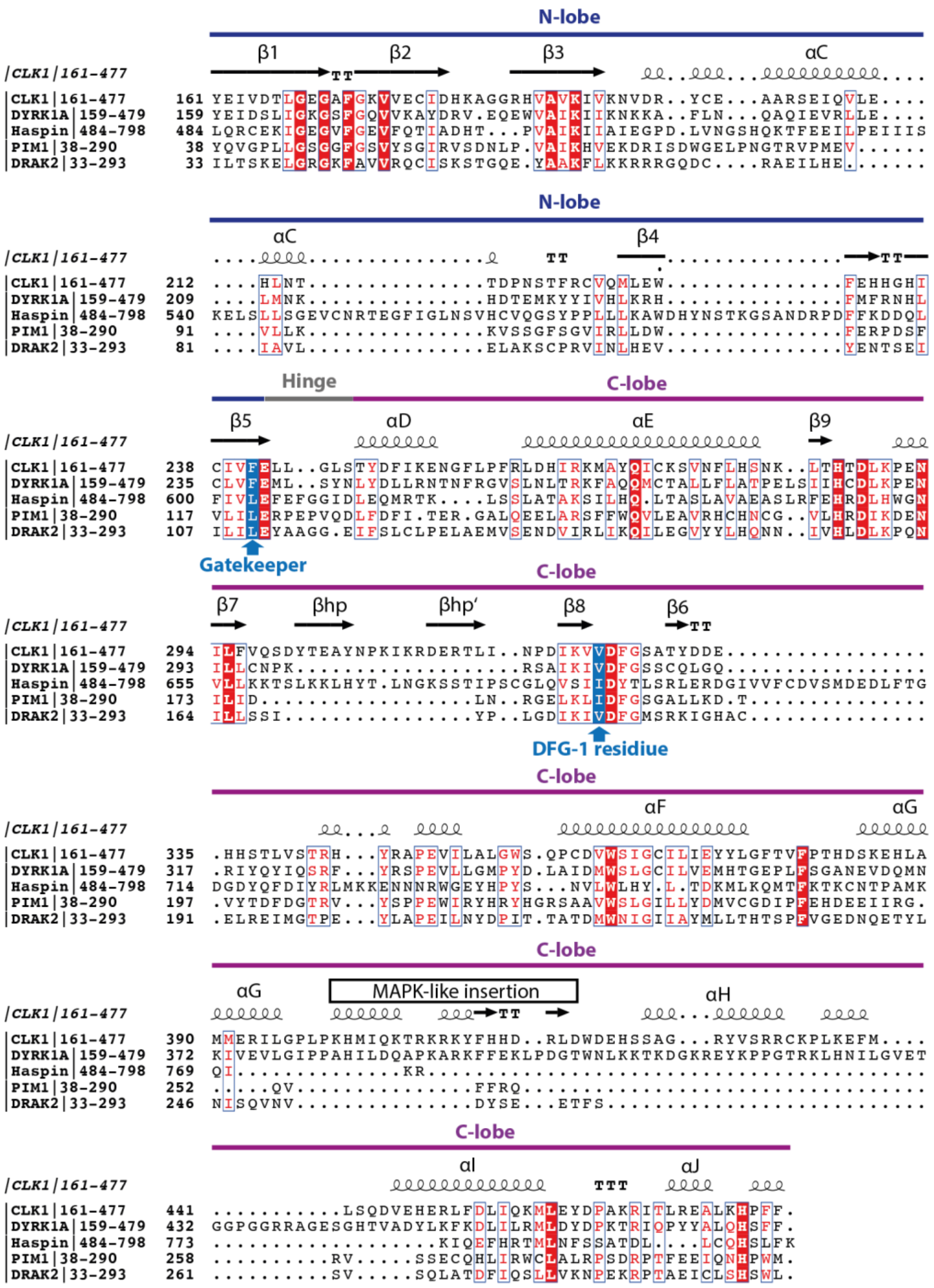


**B**

| Percent Identity | CLK1 161-477 | DYRK1A 159-479 | Haspin 484-798 | PIM1 38-290 | DRAK2 33-293 |
|---|---|---|---|---|---|
| CLK1 161-477 | 100 | | | | |
| DYRK1A 159-479 | 33 | 100 | | | |
| Haspin 484-798 | 17 | 27 | 100 | | |
| PIM1 38-290 | 26 | 24 | 22 | 100 | |
| DRAK2 33-293 | 26 | 26 | 17 | 26 | 100 |

**Supplementary figure 1. Comparative sequence analyses of the most common kinase targets of 1-5.** Sequence alignment of the kinase domains of CLK1, DYRK1A, haspin, PIM1 and DRAK2 is shown in panel **A** and sequence identities in panel **B**.

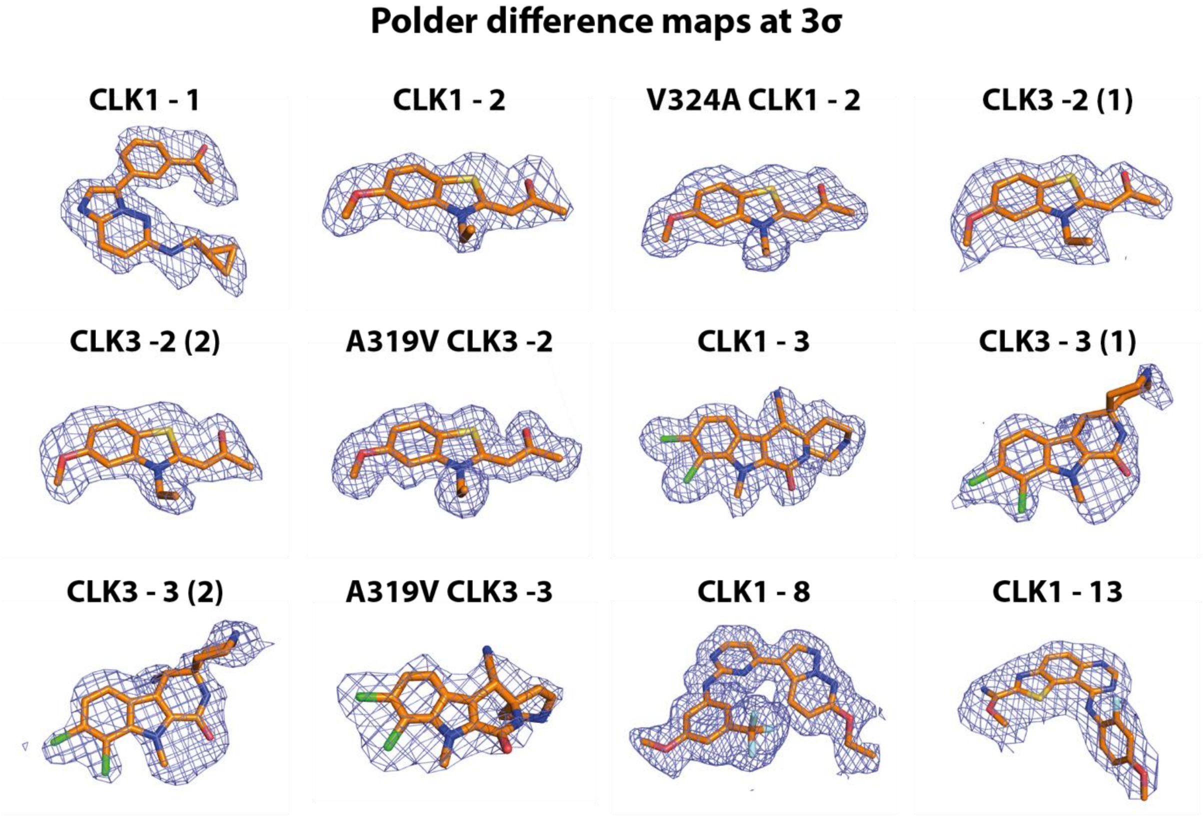


**Supplementary figure 2. Polder difference maps at 3σ for the bound compounds.**

**Supplementary table 1.** Thermal shift results of compounds 1-4 against various kinases.

| | ΔTm (°C) for inhibitor | | | | | | |
|---|---|---|---|---|---|---|---|
| **Kinase** | **1** | **2** | **3** | **4** | **Gatekeeper** | **DFG-1 residue** | **subfamily** |
| AAK1 | 6.9 | 1.5 | 1.5 | 3.4 | **M** | **C** | Other |
| ABL2 | | -0.5 | | | **T** | **A** | TK |
| ACVR2A | 4.0 | | | 0.3 | **T** | **A** | TKL |
| ACVR2B | 7.4 | | | 0.0 | **T** | **A** | TKL |
| AKT3 | 0.4 | 0.0 | 0.4 | | **M** | **T** | AGC |
| ACVRL1 | 11.1 | | | 1.2 | **T** | **A** | TKL |
| ACVR1 | 11.0 | | | 0.7 | **T** | **A** | TKL |
| ACVR1B | 9.1 | | | 1.1 | **S** | **A** | TKL |
| PRKAA1 | 1.3 | -0.3 | | 0.5 | **M** | **A** | CAMK |
| PRKAA2 | 1.1 | -0.1 | | | **M** | **A** | CAMK |
| ADRBK1 | | 0.1 | | | **L** | **S** | AGC |
| ADRBK2 | | -0.2 | 1.1 | | **L** | **S** | AGC |
| BMP2K | 10.7 | 0.2 | 1.0 | | **M** | **C** | Other |
| BMPR1A | 10.9 | | | | **T** | **A** | TKL |
| BMPR1B | 11.5 | | | 0.2 | **T** | **A** | TKL |
| BMPR2 | 7.8 | | 0.3 | | **M** | **S** | TKL |
| BMX | 0.4 | -0.1 | | 0.4 | **T** | **S** | TK |
| PTK6 | 0.4 | | | | **T** | **G** | TK |
| CAMK1D | 0.6 | 0.0 | 1.2 | 0.2 | **M** | **S** | CAMK |
| CAMK1G | 0.6 | -0.1 | | 0.3 | **M** | **T** | CAMK |
| CAMK2A | 0.5 | -0.1 | 1.2 | 0.1 | **F** | **A** | CAMK |
| CAMK2B | 0.2 | 0.3 | | | **F** | **A** | CAMK |
| CAMK2D | | -0.6 | 1.8 | | **F** | **A** | CAMK |
| CAMK2G | | 0.1 | | | **F** | **A** | CAMK |
| CAMK4 | | -0.2 | | | **L** | **A** | CAMK |
| CAMKK1 | | 0.0 | 1.0 | | **F** | **A** | Other |
| CAMKK2 | 2.5 | 0.4 | | | **F** | **A** | Other |
| CDK2 | 1.5 | -0.1 | 0.4 | 0.3 | **F** | **A** | CMGC |
| CDK4 | | -0.6 | | 1.5 | **F** | **A** | CMGC |
| CDK6 | 0.1 | 0.4 | | | **F** | **A** | CMGC |
| CDK8 | 2.7 | 0.1 | 2.2 | 1.9 | **F** | **A** | CMGC |
| CDKL1 | 0.2 | 0.4 | 0.9 | 0.8 | **F** | **C** | CMGC |
| CDKL2 | | -0.5 | | 0.1 | **F** | **C** | CMGC |
| CDKL3 | | 0.1 | | | **F** | **C** | CMGC |
| CDKL5 | | 1.1 | | 0.3 | **F** | **C** | CMGC |
| CHEK2 | 0.8 | 0.1 | 0.3 | 0.5 | **L** | **T** | CAMK |
| CSNK1E | 4.4 | 5.1 | 1.0 | | **M** | **I** | CK1 |
| CSNK1G1 | 1.7 | 3.2 | | 1.0 | **L** | **I** | CK1 |
| CSNK1G2 | 2.5 | 2.7 | | | **L** | **I** | CK1 |
| CSNK1G3 | 0.3 | 3.3 | 2.9 | 1.2 | **L** | **I** | CK1 |
| CSNK2A1 | 3.3 | 2.6 | 3.5 | 0.2 | **F** | **I** | CMGC |
| CSNK2A2 | 3.3 | 2.6 | 4.2 | 0.9 | **F** | **I** | CMGC |
| CLK1 | 5.9 | 6.7 | 7.0 | 7.7 | **F** | **V** | CMGC |
| CLK2 | 5.5 | 5.0 | 3.5 | 7.2 | **F** | **V** | CMGC |
| CLK3 | 1.9 | 2.2 | 2.2 | 1.0 | **F** | **A** | CMGC |
| CLK4 | 8.5 | 9.8 | 4.6 | 0.9 | **F** | **V** | CMGC |
| DAPK3 | 2.8 | 0.9 | 10.6 | | **L** | **I** | CAMK |
| DCLK1 | 0.6 | -0.4 | 0.9 | 0.2 | **M** | **G** | CAMK |
| DDR1 | | 0.4 | | | **T** | **A** | TK |
| DMPK | 1.0 | 1.0 | | 0.8 | **M** | **A** | AGC |
| CDC42BPG | 0.6 | -1.3 | | | **M** | **A** | AGC |
| STK17A | 3.4 | 1.7 | 4.3 | 2.2 | **L** | **V** | CAMK |
| STK17B | 4.2 | 1.0 | 3.4 | 6.9 | **L** | **V** | CAMK |
| DYRK1A | 3.6 | 6.1 | 2.5 | | **F** | **V** | CMGC |

**Supplementary table 1 (cont.).** Thermal shift results of compounds 1-4 against various kinases.

| | ΔTm (°C) for inhibitor | | | | | | |
|---|---|---|---|---|---|---|---|
| **Kinase** | **1** | **2** | **3** | **4** | **Gatekeeper** | **DFG-1 residue** | **subfamily** |
| DYRK2 | 4.1 | 7.3 | 3.6 | 8.9 | F | I | CMGC |
| MAPK3 | 0.5 | -0.1 | 0.5 | | Q | C | CMGC |
| MAPK6 | 0.4 | 0.0 | | | Q | G | CMGC |
| MAPK7 | | 0.2 | | | L | G | CMGC |
| FES | 0.6 | -0.1 | 4.1 | 0.3 | M | S | TK |
| FGFR4 | | | | 0.2 | V | A | TK |
| FGR | | -0.2 | 0.1 | | T | A | TK |
| GAK | 7.3 | 0.2 | 1.4 | 3.3 | T | C | Other |
| GRK5 | | 0.8 | 1.4 | | L | S | AGC |
| GSK3B | 4.1 | 1.4 | 3.9 | | L | C | CMGC |
| GSG2 | 6.5 | | 5.6 | 4.0 | F | I | Other |
| MAP4K4 | | -0.1 | | 0.2 | M | V | STE |
| IGF1R | | | | 0.2 | M | G | TK |
| IKBKB | | 0.1 | 0.6 | 1.5 | M | I | Other |
| ERN2 | | | | 0.4 | L | S | Other |
| ITK | | 1.1 | | | F | S | TK |
| JAK1 | 6.4 | 0.1 | 0.9 | | E | S | TK |
| MAPK8 | | 0.7 | 0.2 | 1.0 | M | L | CMGC |
| MAPK9 | 0.8 | 0.4 | 0.2 | 0.9 | M | L | CMGC |
| LATS1 | | 0.1 | | | M | T | AGC |
| LIMK1 | | | | 0.5 | T | A | TKL |
| STK10 | 3.3 | 0.8 | 0.1 | 1.9 | I | A | STE |
| LYN | 2.4 | 0.1 | 0.2 | 0.3 | T | A | TK |
| MAP2K1 | | -0.1 | | 0.2 | M | C | STE |
| MAP2K2 | 2 | 0.5 | | 0.1 | M | C | STE |
| MAP2K6 | 1.1 | 0.0 | 0.3 | | M | C | STE |
| MAP3K5 | 2.2 | 0.4 | 0.2 | | M | S | STE |
| MERTK | 1.3 | 0.0 | 0.5 | | L | A | TK |
| STK16 | 2.3 | 0.2 | 1.0 | 3.5 | L | M | Other |
| CDC42BPA | | -0.4 | | 0.6 | M | A | AGC |
| CDC42BPB | | 0.3 | | 0.2 | M | A | AGC |
| SRPK3 | 0.4 | 0.0 | 1.9 | 0.2 | L | A | CMGC |
| STK4 | 2.6 | 0.7 | | | M | A | STE |
| STK3 | 2.9 | | 0.3 | 1.6 | M | A | STE |
| STK24 | | -0.2 | | 0.8 | M | A | STE |
| STK26 | 0.3 | -0.3 | | 0.1 | M | A | STE |
| MYO3A | | | | 0.2 | L | V | STE |
| PKMYT1 | 0.9 | -0.6 | | 0.1 | T | G | Other |
| STK38 | 1.1 | 0.4 | 0.4 | | M | S | AGC |
| STK38L | 0.4 | 0.3 | | 0.9 | M | S | AGC |
| NEK1 | | -0.3 | | 0.6 | M | G | Other |
| NEK11 | | 0.6 | | | T | G | Other |
| NEK2 | | 0.8 | | 0.1 | M | G | Other |
| NEK6 | | -1.0 | | | L | G | Other |
| NEK9 | | 0.5 | | | L | G | Other |
| NLK | | -0.2 | | | T | C | CMGC |
| OXSR1 | 0.5 | | | | M | A | STE |
| MAPK11 | 0.3 | 0.0 | 0.2 | | T | L | CMGC |
| MAPK13 | | 0.4 | 0.5 | 0.5 | M | L | CMGC |
| PAK1 | | 0.0 | | | M | T | STE |
| PAK2 | | -0.6 | | | M | T | STE |
| PAK4 | 1.5 | 1.7 | | 1.5 | M | S | STE |
| PAK7 | 0.4 | -0.1 | | 0.6 | M | S | STE |
| PAK6 | 1.1 | -0.9 | | | M | S | STE |
| PBK | | 0.5 | | | M | C | Other |

**Supplementary table 1 (cont.).** Thermal shift results of compounds 1-4 against various kinases.

| | ΔTm (°C) for inhibitor | | | | | | |
|---|---|---|---|---|---|---|---|
| Kinase | 1 | 2 | 3 | 4 | Gatekeeper | DFG-1 residue | subfamily |
| CDK16 | 0.5 | -0.4 | 2.6 | 1.2 | **F** | **A** | CMGC |
| CDK17 | | 0.0 | | 0.4 | **F** | **A** | CMGC |
| PDPK1 | | 0.2 | | 0.3 | **L** | **T** | AGC |
| PHKG2 | 1.7 | 0.2 | | 1.7 | **F** | **S** | CAMK |
| PIM1 | 6.3 | 5.9 | 10.7 | 5.9 | **L** | **I** | CAMK |
| PIM2 | 4.7 | 3.6 | 6.5 | 2.1 | **L** | **I** | CAMK |
| PIM3 | 6.8 | 4.9 | 9.6 | 5.4 | **L** | **I** | CAMK |
| PRKACA | | -0.2 | 3.3 | | **M** | **T** | AGC |
| PRKCZ | | -0.2 | 0.9 | | **I** | **T** | AGC |
| PRKD2 | | 0.7 | 0.9 | | **M** | **C** | CAMK |
| PRKD3 | 0.8 | 2.7 | 2.9 | | **M** | **C** | CAMK |
| PRKG1 | | 1.6 | 6.1 | 0.5 | **M** | **V** | AGC |
| PRKG2 | | -0.1 | | | **L** | **V** | AGC |
| PKN1 | 1.7 | -0.5 | | 0.4 | **M** | **A** | AGC |
| PKN2 | 3.7 | -0.5 | 2.9 | | **M** | **A** | AGC |
| PLK1 | 2.0 | 0.3 | 1.4 | 0.4 | **L** | **G** | Other |
| PLK4 | 2.6 | 0.2 | 6.8 | 0.4 | **L** | **A** | Other |
| PRKX | | -0.2 | 5.8 | | **M** | **T** | AGC |
| SIK2 | | -0.2 | | 0.4 | **T** | **A** | CAMK |
| GRK1 | 1.1 | 0.4 | 2.4 | | **M** | **S** | AGC |
| RIOK2 | 0.3 | -0.2 | | | **M** | **I** | Atypical |
| RIPK2 | 6.0 | | | | **T** | **A** | TKL |
| RIPK3 | | -0.6 | | | **T** | **A** | TKL |
| RPS6KA1 | 5.2 | 0.6 | 1.0 | 0.6 | **L** | **T** | CAMK |
| RPS6KA3 | 3.4 | 0.7 | | | **L** | **T** | CAMK |
| RPS6KA2 | 2.7 | 0.4 | 2.2 | 1.0 | **L** | **T** | CAMK |
| RPS6KA6 | 4.2 | 0.4 | 0.8 | 2.1 | **L** | **T** | CAMK |
| MYLK4 | 5.8 | 0.9 | 3.5 | 3.9 | **M** | **I** | CAMK |
| SgK223 | | | | 0.4 | **T** | **S** | Other |
| SGK3 | | -0.1 | | | **L** | **T** | AGC |
| STK40 | | 0.0 | | | **L** | **T** | CAMK |
| SIK1 | | 1.5 | | | **T** | **A** | CAMK |
| MYLK2 | | -1.4 | | 0.0 | **M** | **I** | CAMK |
| SLK | 2.3 | -0.5 | | 0.5 | **I** | **A** | STE |
| MYLK | | -0.2 | 3.4 | 0.1 | **L** | **I** | CAMK |
| SRPK1 | | 0.0 | | | **F** | **A** | CMGC |
| SRPK2 | | -0.2 | 2.4 | 0.4 | **F** | **A** | CMGC |
| STK33 | 3.1 | 0.6 | 5.9 | 2.4 | **M** | **T** | CAMK |
| STK39 | 0.8 | 0.5 | | 0.0 | **M** | **A** | STE |
| TEC | | 1.0 | | | **T** | **S** | TK |
| TGFBR1 | 9.5 | | | 0.9 | **S** | **A** | TKL |
| TGFBR2 | 13.7 | | | 2.0 | **T** | **C** | TKL |
| TLK1 | | -0.1 | | | **L** | **T** | Other |
| TNIK | 1.2 | 0.1 | | 1.2 | **M** | **V** | STE |
| TRIB1 | | 0.2 | | 0.1 | **L** | **E** | CAMK |
| TTK | 2.7 | 0.5 | 0.7 | 1.7 | **M** | **I** | Other |
| TYK2 | | 0.6 | | | **T** | **S** | TK |
| TYRO3 | 0.4 | 0.1 | | | **L** | **A** | TK |
| VRK1 | 0.7 | 1.1 | 0.3 | | **M** | **V** | CK1 |
| VRK2 | 1.5 | 0.2 | 0.3 | 0.0 | **M** | **A** | CK1 |
| VRK3 | | 0.5 | 0.1 | | **L** | **A** | CK1 |
| WNK3 | 0.0 | -0.3 | | | **T** | **G** | Other |
| STK32A | | -0.2 | 0.8 | 0.2 | **V** | **T** | AGC |
| STK32B | | -0.2 | | | **V** | **T** | AGC |
| STK32C | | -0.9 | 0.7 | | **V** | **T** | AGC |
| STK25 | 0.5 | 0.7 | 1.1 | | **M** | **A** | STE |
| ZAK | 1.4 | -0.3 | | 0.3 | **T** | **C** | TKL |

**Supplementary table 2.** The gatekeeper and DFG-1 amino acid compositions of the kinases that interact with inhibitor **1-5.** The kinases that were test are indicated with T, while those that showed inhibitor binding, either in thermal shift assays or KINOMEscan [1-3], are marked with X. The percentage of the occurrence of each amino acid are shown in Figure 2.

| Kinase | 1 | 2 | 3 | 4 | 5 | Gatekeeper | DFG-1 residue | DFG | subfamily |
|---|---|---|---|---|---|---|---|---|---|
| AAK1 | X | T | T | T |  | M | C | DFG | Other |
| ACVR1 | X | T |  | T |  | T | A | DLG | TKL |
| ACVR1B | X | T |  | T | T | S | A | DLG | TKL |
| ACVR2A | X | T |  | T |  | T | A | DFG | TKL |
| ACVR2B | X | T |  | T |  | T | A | DFG | TKL |
| ACVRL1 | X | T | T | T |  | T | A | DLG | TKL |
| BMP2K | X | T | T |  |  | M | C | DFG | Other |
| BMPR1A | X | T |  |  |  | T | A | DLG | TKL |
| BMPR1B | X | T |  | T |  | T | A | DLG | TKL |
| BMPR2 | X | T | T |  |  | M | S | DFG | TKL |
| CLK1 | X | X | X | X | X | F | V | DFG | CMGC |
| CLK2 | X | X |  | X | X | F | V | DFG | CMGC |
| CLK4 | X | X | X |  | X | F | V | DFG | CMGC |
| CSNK1E | X | X | T |  |  | M | I | DFG | CK1 |
| CSNK1G2 | T | X |  |  | T | L | I | DFG | CK1 |
| CSNK1G3 | T | X | T | T | T | L | I | DFG | CK1 |
| CSNK2A2 | T | T | X | T | T | F | I | DWG | CMGC |
| DAPK3 | T | T | X |  | T | L | I | DFG | CAMK |
| DRAK1 | T | T | X | T | T | L | V | DFG | CAMK |
| DRAK2 | X | T | T | X |  | L | V | DFG | CAMK |
| DYRK1A | T | X | T |  | X | F | V | DFG | CMGC |
| DYRK1B |  | X |  |  | X | F | V | DFG | CMGC |
| DYRK2 | X | X | T | X | X | F | I | DFG | CMGC |
| GAK | X | T | T | T |  | T | C | DFG | Other |
| GSK3A |  | T |  |  | X | L | C | DFG | CMGC |
| GSK3B | X | T | T |  | X | L | C | DFG | CMGC |
| Haspin | X | T | X | X | X | F | I | DYT | Other |
| HIPK2 |  | T |  |  | X | F | I | DFG | CMGC |
| HIPK3 |  | T |  |  | X | F | I | DFG | CMGC |
| IRAK4 |  | T |  |  | X | Y | S | DFG | TKL |
| JAK1 | X | T | T |  | T | E | S | DPG | TK |
| MAP3K19 |  | X |  |  |  | M | I | DFG | STE |
| MYLK4 | X | T | T | T |  | M | I | DFG | CAMK |
| NTRK1 |  | T |  |  | X | F | G | DFG | TK |
| PIM1 | X | X | X | X | X | L | I | DFG | CAMK |
| PIM2 | X | T | X | T | X | L | I | DFG | CAMK |
| PIM3 | X | X | X | X | T | L | I | DFG | CAMK |
| PLK4 | T | T | X | T |  | L | A | DFG | Other |
| PRKG1 |  | T | X | T | T | M | V | DFG | AGC |
| PRKX |  | T | X |  | T | M | T | DFG | AGC |
| RIPK2 | X | T |  |  |  | T | A | DFG | TKL |
| RPS6KA1 | X | T | T | T | T | L | T | DFG | CAMK |
| RPS6KA6 | X | T | T | T | T | L | T | DFG | CAMK |
| STK33 | T | T | X | T | T | M | T | DFG | CAMK |
| TGFbR1 | X | T |  | T | T | S | A | DLG | TKL |
| TGFbR2 | X | T |  | T |  | T | C | DFG | TKL |

**Supplementary table 3.** ΔTm data for wild type and DFG-1-mutated CLK1 and CLK3.

| | | ΔTm (°C) for kinases | | | | |
|---|---|---|---|---|---|---|
| Cpd | name | wild type CLK1 | V324A CLK1 | wild type CLK3 | A319V CLK3 | SMILES |
| 1 | K00135[4] | 7.2 | 4.9 | 1.4 | 5.8 | N(C(=C1)C(=CC2C(=O)C)C=CC=2)(N=C2NCC(C3)C3)C(=N1)C=C2 |
| 2 | Tg003[5] | 7.1 | 3.6 | 1.1 | 5.8 | O=C(C)/C=C1SC2=CC=C(OC)C=C2N\1CC |
| 3 | KH-CARB13[6] | 7.4 | 2.8 | 1.2 | 5.8 | [Cl-].CN1C2C(C(C#N)C3(CC[NH2+]CC3)NC2=O)c4ccc(Cl)c(Cl)c14 |
| 4 | K00972[7] | 5.7 | 2.4 | 0.1 | 0.6 | C(C(C=CC1C(=NC(=NC2)N)C=2)=N2)(C=1)=C(O2)C(=CC=C(C1)[Cl])C=1 |
| 5 | MU1210[2] | 9.3 | 5.94 | 1.5 | 8.27 | Cn1cc(cn1)c2ccc3occ(c4cccc(c4)c5ccncc5)c3n2 |
| 6 | staurosporine | 13.4 | 11.8 | 4.7 | 8.1 | CC12C(C(CC(O1)N3C4=CC=CC=C4C5=C6C(=C7C8=CC=CC=C8N2C7=C53)CNC6=O)NC)OC |
| 7 | KH-CB19[8] | 15.0 | 7.5 | 8.6 | 11 | C(=C1C=C2)(C(=C2[Cl])[Cl])N(C(=C1C(C#N)=CN)C(OCC)=O)C |
| 8 | GW807982X[9] | 9 | 2.5 | 2.0 | 5.5 | CCOc1ccc2c(cnn2n1)c3ccnc(Nc4cc(OC)cc(c4)C(F)(F)F)n3 |
| 9 | K00518 (biofocus) | 6.8 | 3.9 | 1.8 | 8.6 | N(C(=C1)C(=CC2C(=O)C)C=CC=2)(N=C2NC(C(C)C)CO)C(=N1)C=C2 |
| 10 | T3-CLK[10] | 18.7 | 13.7 | 13.7 | 17.3 | CN1CCN(C(C(C)(C)C2=CC=C(C(NC3=CN(C=C(C4=CC=NC=C4)C=C5)C5=N3)=O)C=C2)=O)CC1 |
| 11 | KuWal151[11] | 9.4 | 5.1 | 1.7 | 4.7 | Clc1cccc(c1)c2c[nH]c3c4C(=O)NCc4ccc23 |
| 12 | FC162[12] | 9.5 | 3.0 | 3.6 | 5.5 | O=C1N(C=Nc2ccc3nc(sc3c12)c4cccnc4)C5CC5 |
| 13 | ETH1610[13] | 10.3 | 3.8 | 3.4 | 9.2 | COC(=N)c1nc2ccc3ncnc(Nc4ccc(OC)cc4F)c3c2s1 |
| 14 | VN412[2] | 12.3 | 7.8 | 4.8 | 10.1 | Cn1cc(cn1)c2ccc3occ(c4cccc(Oc5ccccc5)c4)c3n2 |
| 15 | GW779439X[9] | 15.0 | 9.8 | 5.7 | 10.2 | CN1CCN(CC1)C2=C(C=C(C=C2)NC3=NC=CC(=N3)C4=C5C=CC=NN5N=C4)C(F)(F)F |
| 16 | KH-CARB10[6] | 6.1 | 1.8 | 1.1 | 5.0 | CN1CCC2(CC1)NC(=O)C3C(C2C#N)c4ccc(Cl)c(Cl)c4N3C |
| 17 | KH-CARB11[6] | 4.8 | 1.0 | 0.6 | 3.8 | CCN1CCC2(CC1)NC(=O)C3C(C2C#N)c4ccc(Cl)c(Cl)c4N3C |
| 18 | iodotubercidin | 13.1 | 9.7 | 7.9 | 14.1 | C1=C(C2=C(N=CN=C2N1C3C(C(C(O3)CO)O)O)N)I |

**Supplementary table 4.** Inhibition constant ($K_i$) from nanoBRET assays for CLK1 wild type and V324A mutant.

| | Ki (µM) | | |
|---|---|---|---|
| **Compound** | **wild type CLK1** | **V324A CLK1** | **ratio mutant/wild type** |
| 1 | 0.229 ± 0.10 | 0.512 ± 0.10 | 2.2 |
| 2 | 0.21 ± 0.12 | 0.671 ± 0.23 | 3.2 |
| 3 | 0.136 ± 0.09 | 2.81 ± 0.67 | 20.6 |
| 6 | 0.009 ± 0.002 | 0.006 ± 0.002 | 0.7 |
| 7 | 0.018 ± 0.008 | 0.166 ± 0.031 | 9.2 |
| 8 | 0.036 ± 0.02 | 3.93 ± 1.1 | 109.2 |
| 9 | 0.943 ± 0.64 | 2.64 ± 1.80 | 2.8 |
| 10 | 0.001 ± 0.0001 | 0.0023 ± 0.0001 | 2.4 |
| 11 | 0.228 ± 0.08 | 0.878 ± 0.046 | 3.9 |
| 12 | 1.86 ± 1.2 | 5.45 ± 0.79 | 2.9 |

**Supplementary table 5.** Data collection and refinement statistics.

| Complex | CLK1-1 | CLK1-2 | V324A CLK1-2 | CLK3-2 | A319V CLK3-2 |
|---|---|---|---|---|---|
| **PDB accession code** | **6YTA** | **6YTE** | **6YTD** | **6YTW** | **6YTY** |
| **Beamline** | **SLS PXIII-X06DA** | **BESSY 14.2** | **BESSY 14.2** | **BESSY 14.2** | **BESSY 14.2** |
| ***Data Collection*** | | | | | |
| Resolution[a] (Å) | 67.46-1.95 (2.00-1.95) | 64.70-2.30 (2.38-2.30) | 71.30-2.00 (2.05-2.00) | 79.52-2.00 (2.05-2.00) | 53.10-1.76 (1.80-1.76) |
| Spacegroup | *C*2 | *C*2 | *I*2 | *C*2 | *I*2 |
| Cell dimensions | $a$ = 92.9, $b$ = 64.2, $c$ = 80.8 Å<br>$\alpha = \gamma$ = 90.0˚; $\beta$ = 123.4˚ | $a$ = 91.7, $b$ = 64.3, $c$ = 73.1 Å<br>$\alpha = \gamma$ = 90.0˚; $\beta$ = 117.7˚ | $a$ = 80.9, $b$ = 64.7, $c$ = 89.3 Å<br>$\alpha = \gamma$ = 90.0˚; $\beta$ = 114.7˚ | $a$ = 96.1, $b$ = 131.6, $c$ = 83.6 Å<br>$\alpha = \gamma$ = 90.0˚; $\beta$ = 108.0˚ | $a$ = 84.2, $b$ = 45.0, $c$ = 106.3 Å<br>$\alpha = \gamma$ = 90.0˚; $\beta$ = 111.1˚ |
| No. unique reflections[a] | 29,054 (2,009) | 16,851 (1,648) | 27,570 (2,045) | 66,435 (4,475) | 36,800 (2,104) |
| Completeness[a] (%) | 100.0 (100) | 100.0 (100.0) | 97.1 (98.0) | 99.8 (100.0) | 99.9 (99.8) |
| I/σI[a] | 12.8 (3.7) | 8.0 (3.7) | 10.2 (2.8) | 11.1 (2.8) | 10.7 (3.1) |
| $R_{merge}$[a] | 0.077 (0.413) | 0.131 (0.383) | 0.097 (0.618) | 0.104 (0.653) | 0.097 (0.553) |
| CC (1/2) | 0.998 (0.909) | 0.969 (0.893) | 0.996 (0.851) | 0.998 (0.839) | 0.995 (0.793) |
| Redundancy[a] | 6.3 (5.9) | 4.8 (4.9) | 6.0 (6.2) | 6.3 (6.7) | 5.2 (5.3) |
| ***Refinement*** | | | | | |
| No. atoms in refinement (P/L/O)[b] | 2,716/23/220 | 2,810/ 17/ 233 | 2,779/ 17/ 254 | 5,686/ 34/ 623 | 2,912/ 17/ 186 |
| B factor (P/L/O)[b] (Å$^2$) | 24/23/26 | 26/ 16/ 29 | 31/ 20/ 39 | 36/ 52/ 40 | 18/ 13/ 20 |
| $R_{fact}$ (%) | 19.0 | 18.1 | 17.9 | 18.4 | 18.6 |
| $R_{free}$ (%) | 27.5 | 25.8 | 22.8 | 22.7 | 22.2 |
| rms deviation bond[c] (Å) | 0.013 | 0.013 | 0.013 | 0.012 | 0.014 |
| rms deviation angle[c] (°) | 1.9 | 1.6 | 1.6 | 1.7 | 1.7 |
| ***Molprobity Ramachandran*** | | | | | |
| Favour (%) | 94.05 | 94.05 | 95.25 | 95.00 | 96.56 |
| Outlier (%) | 0 | 0 | 0 | 0.15 | 0 |
| Crystallization conditions | 14% PEG 6k, 0.1M bicine 8.0 | 26% PEG 6k, 0.1M bicine 9.0 | 17% PEG 3350, 0,2M Na malonate pH 7 | 21% PEG 3350, 0,2M Na/K PO4, 10% Ethylene Glycol | 17% PEG 3350, 0,2M NaBr, 10% Ethylene Glycol, 0.1M bis-tris propane 7.0 |

[a] Values in brackets show the statistics for the highest resolution shells.

[b] P/L/O indicate protein, ligand molecules presented in the active sites, and other (water and solvent molecules), respectively.

[c] rms indicates root-mean-square.

**Supplementary table 5 (cont.).** Data collection and refinement statistics.

| **Complex** | **CLK1-3** | **CLK3-3** | **A319V CLK3-3** | **CLK1-8** | **CLK1-13** |
|---|---|---|---|---|---|
| **PDB accession code** | **6YTG** | **6YU1** | **6Z2V** | **6ZLN** | **6YTI** |
| **Beamline** | **SLS PXIII-X06DA** | **BESSY 14.1** | **BESSY 14.2** | **SLS PXI-X06SA** | **SLS PXIII-X06DA** |
| ***Data Collection*** | | | | | |
| Resolution[a] (Å) | 64.30-1.95 (2.00-1.95) | 79.66-1.90 (1.94-1.90) | 76.54-2.60 (2.72-2.60) | 45.53-1.70 (1.73-1.70) | 69.57-2.40 (2.49-2.40) |
| Spacegroup | *C*2 | *C*2 | *I*2 | *C*2 | *C*2 |
| Cell dimensions | $a$ = 92.49, $b$ = 64.1, $c$ = 80.8 Å<br>$\alpha$ = $\gamma$ = 90.0˚; $\beta$ = 123.4˚ | $a$ = 96.3, $b$ = 131.0, $c$ = 83.6 Å<br>$\alpha$ = $\gamma$ = 90.0˚; $\beta$ = 107.7˚ | $a$ = 84.4, $b$ = 45.4, $c$ = 106.3 Å<br>$\alpha$ = $\gamma$ = 90.0˚; $\beta$ = 111.2˚ | $a$ = 91.6, $b$ = 63.6, $c$ = 80.1 Å<br>$\alpha$ = $\gamma$ = 90.0˚; $\beta$ = 118.4˚ | $a$ = 91.5, $b$ = 63.9, $c$ = 79.4 Å<br>$\alpha$ = $\gamma$ = 90.0˚; $\beta$ = 118.8˚ |
| No. unique reflections[a] | 29,054 (2,009) | 76,479 (4,232) | 11,812 (1,441) | 44,218 (2,200) | 15,727 (1,633) |
| Completeness[a] (%) | 100 (100) | 98.7 (91.8) | 100.0 (100.0) | 99.42(93.8) | 99.4 (98.9) |
| I/σI[a] | 26.8 (3.7) | 10.7 (2.7) | 6.4 (1.6) | 11.1 (2.4) | 10.1 (1.9) |
| $R_{merge}$[a] | 0.071 (0.374) | 0.073 (0.402) | 0.246 (1.319) | 0.065 (0.417) | 0.153 (1.064) |
| CC (1/2) | 0.998 (0.909) | 0.997 (0.838) | 0.985 (0.597) | 0.997 (0.839) | 0.996(0.707 |
| Redundancy[a] | 6.3 (5.9) | 4.0 (3.4) | 6.9 (7.2) | 3.9 (3.6) | 7.0 (6.9) |
| ***Refinement*** | | | | | |
| No. atoms in refinement (P/L/O)[b] | 2,685/48/233 | 5,707/48/579 | 2,761/ 24/ 66 | 2,783/62/415 | 2,665/27/103 |
| B factor (P/L/O)[b] (Å$^2$) | 29/24/32 | 29/60/40 | 39/ 61/ 29 | 23/21/35 | 49/48/46 |
| $R_{fact}$ (%) | 20.4 | 17.1 | 20.2 | 17.0 | 19.5 |
| $R_{free}$ (%) | 27.0 | 20.7 | 26.4 | 19.9 | 25.6 |
| rms deviation bond[c] (Å) | 0.015 | 0.011 | 0.017 | 0.015 | 0.012 |
| rms deviation angle[c] (°) | 1.7 | 1.6 | 1.8 | 1.7 | 1.7 |
| ***Molprobity Ramachandran*** | | | | | |
| Favour (%) | 94.20 | 96.04 | 91.04 | 96.73 | 94.12 |
| Outlier (%) | 0 | 0.15 | 0.30 | 0.30 | 0.31 |
| Crystallization conditions | 17% PEG 6k, 0.1M bicine 8.0 | 18% PEG 3350, 0,2M Na/K PO4, 10% Ethylene Glycol - - | 24% PEG 3350, 0,2M KSCN, 10% Ethylene Glycol, 0.1M bis-tris propane 6.5 | 14% PEG 6k, 0.1M bicine 9.0 | 29% PEG 6k, 0.1M bicine 9.3 |

[a] Values in brackets show the statistics for the highest resolution shells.

[b] P/L/O indicate protein, ligand molecules presented in the active sites, and other (water and solvent molecules), respectively.

[c] rms indicates root-mean-square.

**Supplementary table 5 (cont.).** Data collection and refinement statistics.

| **Complex** | **ACVR1-1** | **CLK2-Ro-3306** |
|---|---|---|
| **PDB accession code** | **4DYM** | **3NR9** |
| **Beamline** | **Diamond I02** | **Diamond I24** |
| ***Data Collection*** | | |
| Resolution[a] (Å) | 44.73-2.42 (2.55-2.42) | 55.90-2.89 (3.04-2.89) |
| Spacegroup | $C222_1$ | $P322_1$ |
| Cell dimensions | $a$ = 57.8, $b$ = 81.86, $c$ = 140.39 Å<br>$\alpha = \gamma = \beta$ = 90.0˚ | $a$ = b = 97.7, $c$ = 223.0 Å<br>$\alpha = \gamma$ = 90.0˚; $\beta$ = 120˚ |
| No. unique reflections[a] | 13,074 (1,858) | 28,133 (4,062) |
| Completeness[a] (%) | 99.8 (100.0) | 99.3 (99.2) |
| I/σI[a] | 8.1 (2.0) | 8.5 (2.0) |
| $R_{merge}$[a] | 0.146 (0.75) | 0.173 (0.989) |
| CC (1/2) | | |
| Redundancy[a] | 4.5 (4.7) | 4.9 (5.0) |
| ***Refinement*** | | |
| No. atoms in refinement (P/L/O)[b] | 2,306/23/158 | 8,427/72/63 |
| B factor (P/L/O)[b] (Å$^2$) | 43/29/40 | 45/39/26 |
| $R_{fact}$ (%) | 22.0 | 19.4 |
| $R_{free}$ (%) | 28.0 | 25.2 |
| rms deviation bond[c] (Å) | 0.012 | 0.013 |
| rms deviation angle[c] (°) | 1.5 | 1.4 |
| ***Molprobity Ramachandran*** | | |
| Favour (%) | 96.91 | 95.13 |
| Outlier (%) | 0.69 | 0.19 |
| Crystallization conditions | 1.60M MgSO4; 0.1M MES pH 6.5 | 1.60M MgSO4; 0.1M MES pH 6.5 |

[a] Values in brackets show the statistics for the highest resolution shells.

[b] P/L/O indicate protein, ligand molecules presented in the active sites, and other (water and solvent molecules), respectively.

[c] rms indicates root-mean-square.
